# Decadal warming is linked to thermal tolerance and transcriptome reprogramming in a seagrass

**DOI:** 10.64898/2026.07.26.740759

**Authors:** Hung Manh Nguyen, Neta Ly Lipkin, Moran Kaminer, Gidon Winters, Simon Barak

**Author notes:** Corresponding author. (S.B.).

## Abstract

Ocean warming and marine heatwaves threaten seagrasses, which are considered highly vulnerable to rising temperatures. We compared thermal stress responses of the tropical seagrass *Halophila stipulacea* collected from the Gulf of Aqaba in 2017 and 2022. Under mesocosm-applied thermal stress, 2017 plants showed reduced growth, impaired photochemistry, and extensive transcriptomic changes. In contrast, 2022 plants maintained normal growth and photochemistry with minimal transcriptional responses. The enhanced thermal stress tolerance of 2022 plants was associated with a “stress-ready” transcriptome, with many stress-response genes already up/downregulated under control conditions, including those encoding heat, oxidative stress, and abscisic acid signaling proteins. These changes coincided with local ocean warming and a shift from episodic to chronic thermal stress. Our findings provide the first longitudinal dataset in a seagrass that tracks physiological and molecular changes occurring concurrently with documented ocean warming.

## INTRODUCTION

Climate change is reshaping terrestrial and marine plant communities globally (*1, 2*). Long-term monitoring and meta-analyses have revealed that warming drives profound changes in species composition including substantial extinction of native plants (*3-5*). In marine ecosystems, the impact of ocean warming is further compounded by the increase in frequency, intensity, and duration of marine heatwaves across most regions (*2, 6*). These thermal extremes disproportionately affect marine foundation species, whose structural and functional roles underpin coastal ecosystems and whose responses can cascade across entire food webs (*7*). For instance, foundation species such as brown macroalgae and seagrasses are projected to experience dramatic contractions in high-quality habitat (78-96% globally) by 2100 under intermediate-to-high emission scenarios, even when net losses in local diversity appear more modest (3-7%) (*7, 8*).

Seagrasses are marine flowering plants distributed along shallow-water coastlines worldwide and provide a wide range of invaluable ecosystem services that support biodiversity, carbon sequestration, and coastal protection (*9, 10*). Global assessments indicate that seagrass meadows have experienced persistent net losses over the past century, with early syntheses estimating declines of ∼110 km^2^ yr^−1^ since the 1980s and accelerated losses in the late 20th century (*11*). More recent global reconstructions confirm substantial long-term area loss (∼19% of surveyed meadow area since the late 1800s), while emphasizing strong regional variability, partial stabilization or recovery in some bioregions, and continued uncertainty due to incomplete global monitoring (*10, 12*).

Ocean warming and increasing occurrences of marine heatwaves are emerging as dominant drivers of seagrass vulnerability, with comprehensive reviews indicating that thermal extremes are likely to accelerate habitat degradation, reduce biomass, and impair key ecosystem services under continued warming (*13, 14*). Global-scale projections further suggest average reductions in above-ground biomass of 4-9% by 2100, with the most severe contractions occurring in both tropical and temperate regions (*15*). These regional changes will be accompanied by substantial loss of suitable habitat at low-latitude range edges, regime shifts to turf-dominated or bare substrates and pronounced reshuffling of evolutionary lineages across regions (*16-18*).

Despite these widespread losses, increasing evidence indicates that physiological and molecular adjustment can buffer some seagrass populations against escalating thermal stress. Responses to marine heatwaves depend strongly on event intensity, duration, and recurrence; extreme episodes often cause widespread mortality, whereas repeated moderate exposures can promote recovery and enhance tolerance (*6, 7*). Comparative studies of stress-tolerant versus stress-sensitive seagrass populations have revealed physiological and transcriptomic signatures of local adjustment and thermal resilience (*19-23*). However, evidence for these signatures has been derived via space-for-time substitution comparisons among populations rather than from repeated observations on the same populations over time. Consequently, direct empirical links between transcriptome reprogramming and long-term adjustment to ocean warming are largely absent due to the lack of longitudinal, multi-year transcriptomic datasets capable of tracking adaptive trajectories over time.

Here, we show evidence for a rapid increase in thermal tolerance in natural populations of the tropical seagrass *Halophila stipulacea* within a single decade. Plants collected from the Gulf of Aqaba in 2022 displayed no reduction in growth or photochemistry under experimental warming, unlike conspecifics collected in 2017. This enhanced thermal tolerance was linked to the acquisition of a “stress-ready” transcriptome in which thousands of stress-responsive genes in the 2017 cohort were already constitutively expressed in the 2022 plants under control conditions. These genes exhibited baseline expression levels prior to stress that were similar to the post-stress levels observed in the 2017 conspecifics, a shift that coincided with local ocean warming and a sharp rise in extreme-temperature days in only five years. Our findings demonstrate that contemporary ocean warming can drive rapid physiological and transcriptomic adjustment in seagrasses over ecologically relevant timescales. More broadly, this rapid stabilization of a protective molecular state provides critical field evidence that wild plant populations can adjust to rising temperatures over multi-year timeframes.

## RESULTS

### Adjustment of *Halophila stipulacea* populations over time to ocean warming and marine heatwaves

Our previous work examining the effect of single and combined thermal and nutrient stresses had revealed little impact of thermal stress on the growth of two populations of mesocosm-grown *H. stipulacea* collected in 2019 from the northern Gulf of Aqaba (GoA; *22, 24*). To confirm this observation, we collected *H. stipulacea* plants from two different sites in the northern GoA in 2022. The North Beach (NB) site adjacent to the city of Eilat represents an anthropogenically-impacted site, whereas the South Beach (SB) site, located away from urban activities, is considered relatively pristine (Fig. 1, A and B; *25-27*). Plants were placed in aquaria and exposed to either 26 °C (control) or ecologically relevant thermal stress of 32 °C (Fig. 1, C to D; see materials and methods). The growth and photochemistry of plants from both sites were not significantly altered even after 35 days of continuous thermal stress (Fig. 1, E and F; Supplementary Table S1), thereby confirming our prior observation that plants collected in 2019 were not affected by elevated temperature (*22, 24*). The tolerance to thermal stress of *H. stipulacea* plants in 2019 and 2022 was in sharp contrast to the effect of thermal stress on conspecifics collected from NB in 2017, where plant growth and photochemistry were significantly reduced by 60% and 20%, respectively, after 21 days at 32 °C (Fig. 1, E and F; Supplementary Table S1; *28*).

**Fig. 1.**
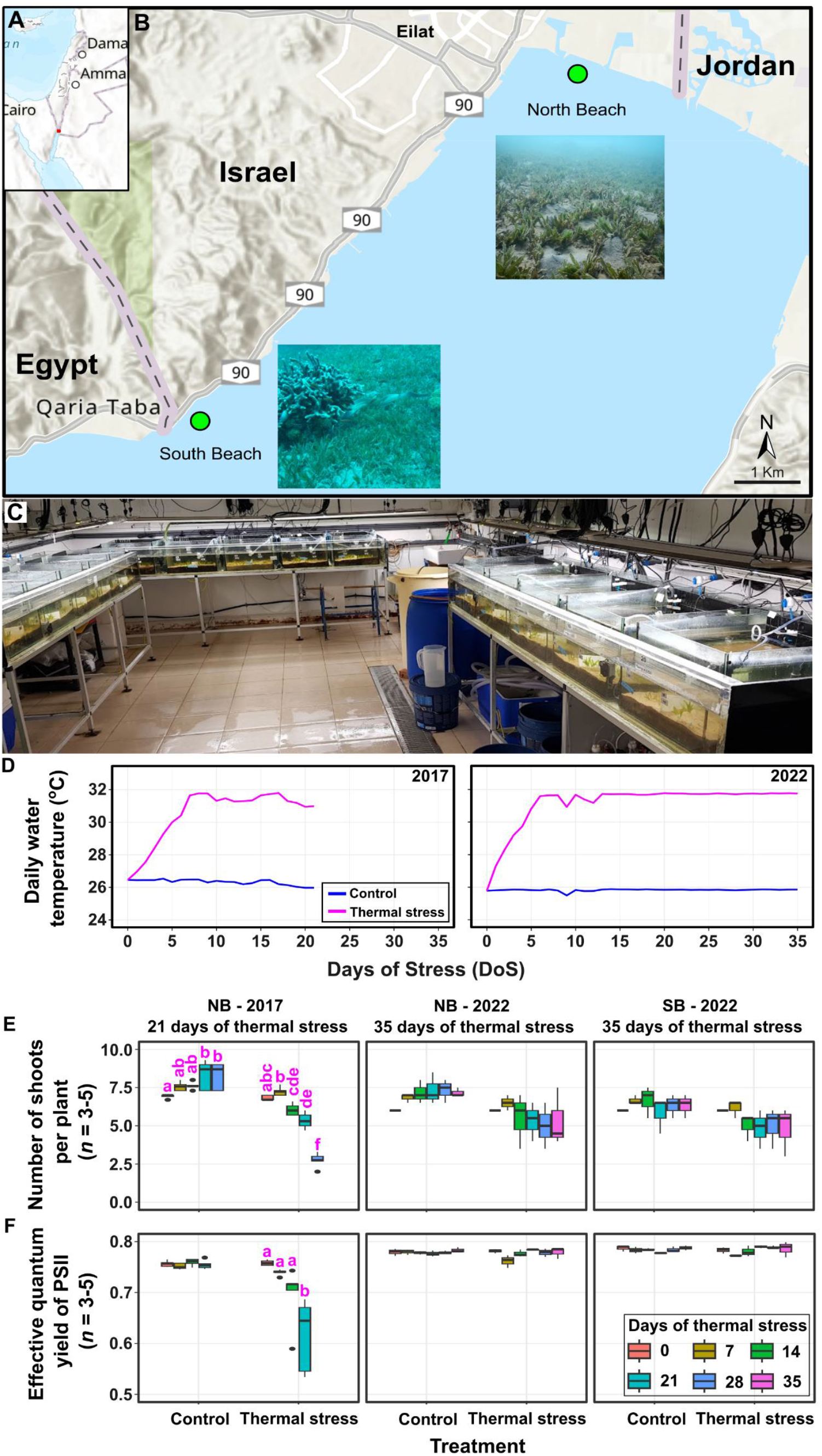
*Halophila stipulacea* sample collection sites and mesocosm experiments. (**A** and **B**) Seagrass plants were collected from the impacted North Beach and pristine South Beach sites (green dots) along the Israeli coastline in the Gulf of Aqaba. Insert photographs show *H. stipulacea* meadows at the North Beach and South Beach sites. Photo credits: Hung Manh Nguyen and Jonas Reghelin de Azevedo. (**C**) The mesocosm system, Dead Sea and Arava Science Center, Israel. (**D**) Temperature profiles of the 2017 and 2022 experiments. (**E** and **F**) Morphological and photo-physiological responses of 2017 and 2022 plants under control and thermal stress treatments. For box and whisker plots, the median (thick black line) and interquartile range are shown. Whiskers indicate the maximum/minimum range. Data were evaluated using a two-way ANOVA (Supplementary Table S1). Prior to analysis, the homogeneity of variance assumption was checked using Levene’s test. Normality was assessed using the Shapiro-Wilk test. Letters indicate significant differences (*p* < 0.05; Tukey HSD *post hoc* test, *n* = 3 to 5) between different days of stress within each treatment.

The increased thermal tolerance of *H. stipulacea* plants in 2019 and 2022 compared to 2017 suggested adjustment to changing environmental conditions. To evaluate whether these phenotypic shifts coincided with local ocean warming, we analyzed long-term seawater temperature records from the Israel National Monitoring Program in the Gulf of Eilat (Gulf of Aqaba; GoA; www.meteo-tech.co.il). Average Maximum Daily Seawater Temperature (MDST) in the GoA increased approximately 0.57 °C between 2006 and 2022, and by 0.66 °C between 2017 and 2022, the period of the current study (Fig. 2A; see materials and methods). Strikingly, the annual number of days exceeding an MDST of 29 °C increased sharply from 1-5 days yr^−1^ between 2013 and 2016 to 13-22 days yr^−1^ between 2018 and 2021 (Fig. 2B) indicating a transition from episodic to chronic thermal stress (*29*). These results suggest a link between progressive seawater warming and enhanced thermal tolerance in the seagrass.

**Fig. 2.**
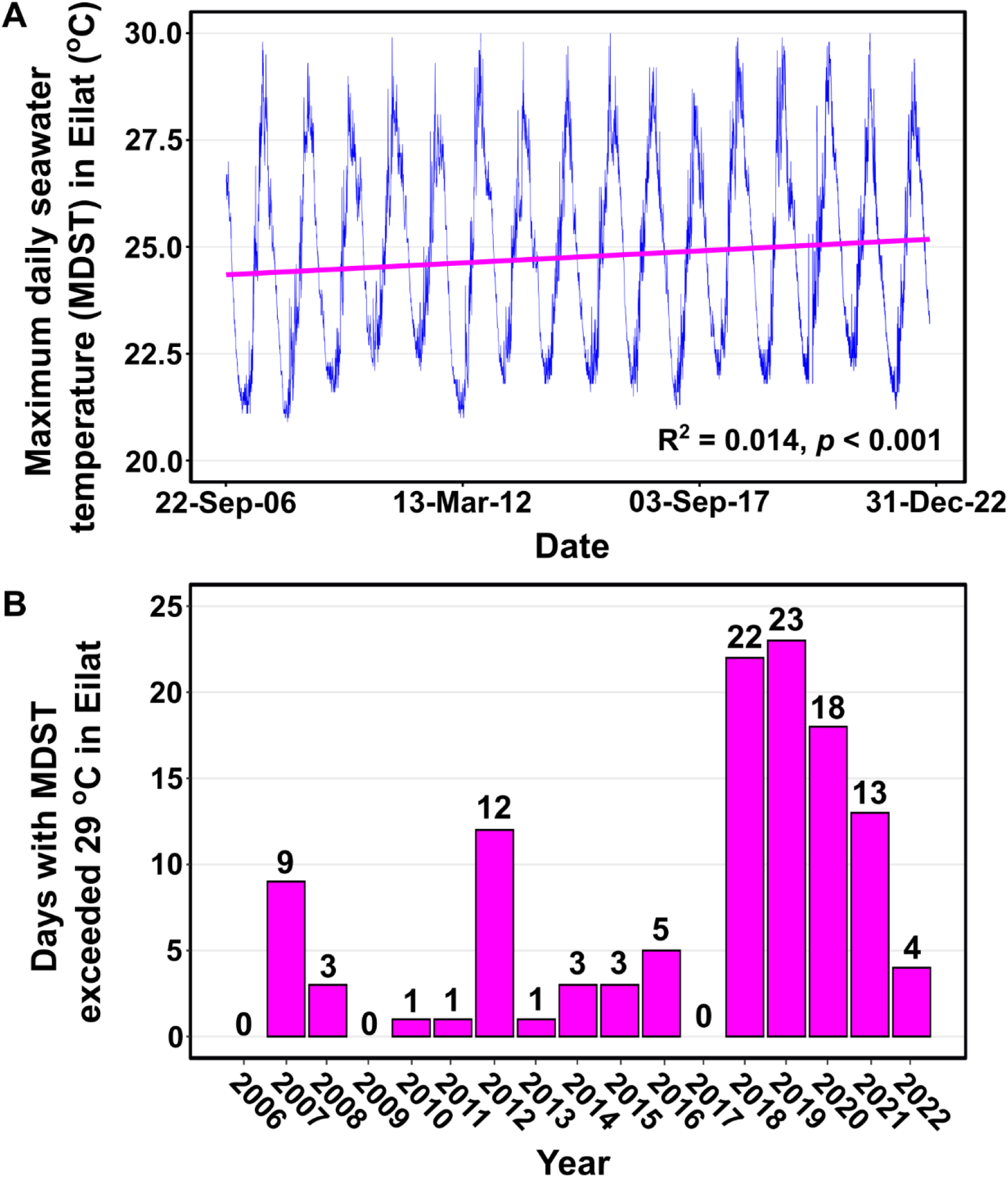
Long-term monitoring data of seawater temperature in the Gulf of Aqaba, Israel between 2006 and 2022. (**A**) Maximum Daily Seawater Temperature (MDST; blue line); Linear regression slope (magenta line), R^2^ and *p*-values from a linear regression model. (**B**) Number of days when the MDST exceeded 29 °C in Eilat. Data were collected from the Israel National Monitoring Program in the Gulf of Eilat (Gulf of Aqaba; GoA; www.meteo-tech.co.il).

### Five-year temporal divergence in *Halophila stipulacea* thermal stress transcriptomes

We performed RNA-seq-based transcript profiling on RNA from *H. stipulacea* plants collected in 2017 (NB) and 2022 (NB and SB) (Supplementary Table S2). These plants had been exposed to the same mesocosm-applied control and thermal stress conditions (21 days for 2017 plants and 35 days for 2022 plants; Fig. 1D; *28*). This provided a unique opportunity to compare thermal stress response transcriptomes between cohorts separated by five years of *in situ* thermal history. RNA quality metrics and the analysis of read lengths and the proportion of non-genic reads showed no evidence of reduced RNA integrity in the 2017 samples, despite their longer storage time prior to sequencing (Supplementary Table S3; Supplementary Fig. S1).

Principal component analysis of global transcriptome adjustment revealed distinct transcriptional reprogramming in response to thermal stress in plants from 2017, whereas control and thermal stress transcriptome samples from 2022 plants clustered together indicating minimal thermal stress-induced transcriptome adjustment (Fig. 3A). The observation that thermal stress had little effect on the transcriptome of 2022 plants compared to 2017 plants, was further investigated by comparing the number of differentially expressed genes (DEGs) in response to thermal stress, between each population. Plants collected from NB in 2017 and exposed to thermal stress in the mesocosms, exhibited 5,095 DEGs (2,548, upregulated; 2,547, downregulated) (Fig. 3B, Supplementary Table S4). However, plants collected from both NB and SB in 2022 and exposed to thermal stress showed comparatively few DEGs (NB: 41, upregulated; 66, downregulated [107 total DEGs]; SB: 89, upregulated; 186, downregulated [275 total DEGs]. GO-term overrepresentation analysis showed that thermal stress-mediated upregulated DEGs in the 2017 plants were enriched in GO-terms linked to the response to stress including: osmotic and salt stresses, the response to abscisic acid, the response to hypoxia and the regulation of response to stress (Fig. 3C, top left panel: Fig. 3D, Clusters 1 and 3; Supplementary Table S5). DEGs that were downregulated by thermal stress were also enriched in GO-terms for processes related to stress including photosynthesis, cell wall organization and sexual reproduction (Fig. 3C, top right panel; Fig. 3D; Clusters 1, 3 and 4; Supplementary Table S5). Thermal stress usually leads to a reduction in photosynthesis, to cell wall remodelling and to deleterious effects on the development of sexual reproductive organs (*30-32*). Conversely, no GO-terms related to stress were enriched in upregulated or downregulated DEGs in 2022 plants from either NB or SB (Fig. 3C, middle and bottom panels; Supplementary Table S5). These results suggest that stress responses were only activated *de novo* in 2017 plants exposed to thermal stress.

**Fig. 3.**
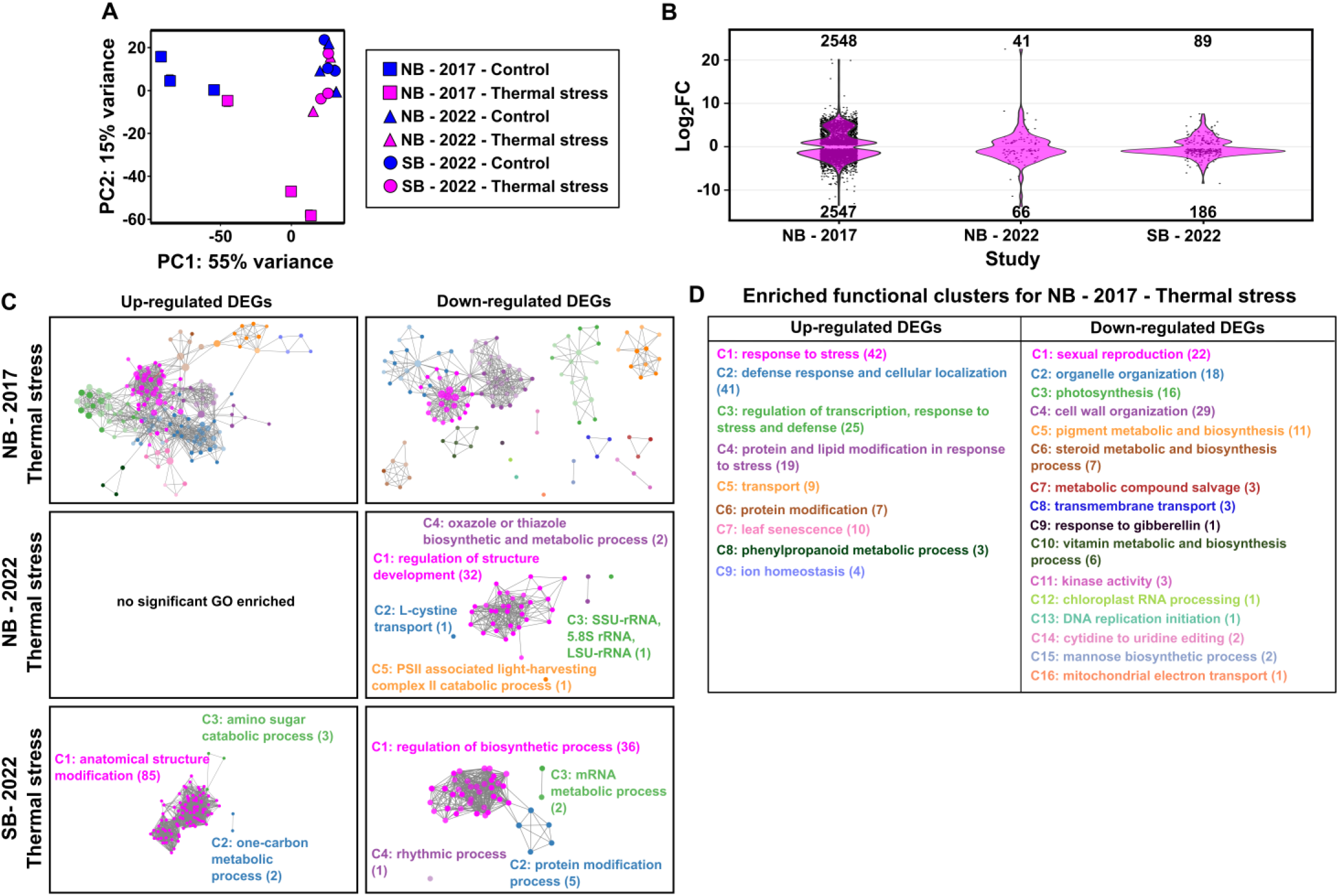
Global transcriptome changes of 2017 and 2022 *H. stipulacea* plants in response to thermal stress. (**A**) Principal component analysis of transcript profiles. Each point represents one biological replicate and each condition is filled with the same-colored symbol while different populations/years are depicted with different symbols (see legend box). (**B**) Log_2_ fold-change of DEGs from 2017 and 2022 plants in response to thermal stress. Numbers above/below plots, total number of up-/downregulated DEGs, respectively. (**C**) Functional clusters enriched for up- and downregulated DEGs in response to thermal stress. Each colored cluster generated by GOMCL (*57*) is labelled with the representative functional term. Each node represents a GO term. Node size, the number of genes in the test set assigned to that functional term; the number of genes in each cluster is in parentheses. Node shade, *p*-value (false discovery rate (FDR)- adjusted *p* < 0.05, *n* = 3). Darker shades indicate smaller *p*-values. GO-terms sharing > 50% of genes are connected by edges. (**D**) Enriched functional clusters for plants collected in 2017 North Beach (NB) and exposed to thermal stress.

The attenuated transcriptional responsiveness to stress of 2022 plants, mirrors “stress-ready” transcriptomes observed in halophytic Brassicaceae. In these plants, genes that are induced or repressed by ionic stress in stress-sensitive *Arabidopsis thaliana* show constitutively high or low expression, respectively, even under control (stress-neutral) conditions (*33-35*). We therefore examined whether the 2022 plants exhibited signatures of a “stress-ready” transcriptome by assigning DEGs from plants from both years to four idealized transcriptional response modes (Fig. 4A; Supplementary Table S6): “Stress-ready”, “Shared response” (where expression of genes is up-/downregulated in plants from both years), “Unique response” (where expression of genes exhibits a response to thermal stress specifically in plants of one year but not in the other), and “Opposite response” (where expression of genes in plants from one year shows the opposite response to plants from the other year). Of the genes categorized within the four response modes, 50.3% and 44.7% of the DEGs in the 2022 NB and SB populations, respectively, displayed a “Stress-ready” mode whereas only 0.7% and 1.9% of DEGs in 2017 plants relative to the 2022 NB and SB plants, respectively, showed a “Stress-ready” mode. In contrast, the largest proportion of DEGs in the 2017 plants exhibited a “Unique response” (47.7% and 50.1% relative to the 2022 NB and SB plants, respectively) while only 0.8% and 1.8% of 2022 NB and SB DEGs showed a “Unique Response”. Fewer than 1% of genes displayed “Shared response” or “Opposite response” modes in plants from either year.

**Fig. 4.**
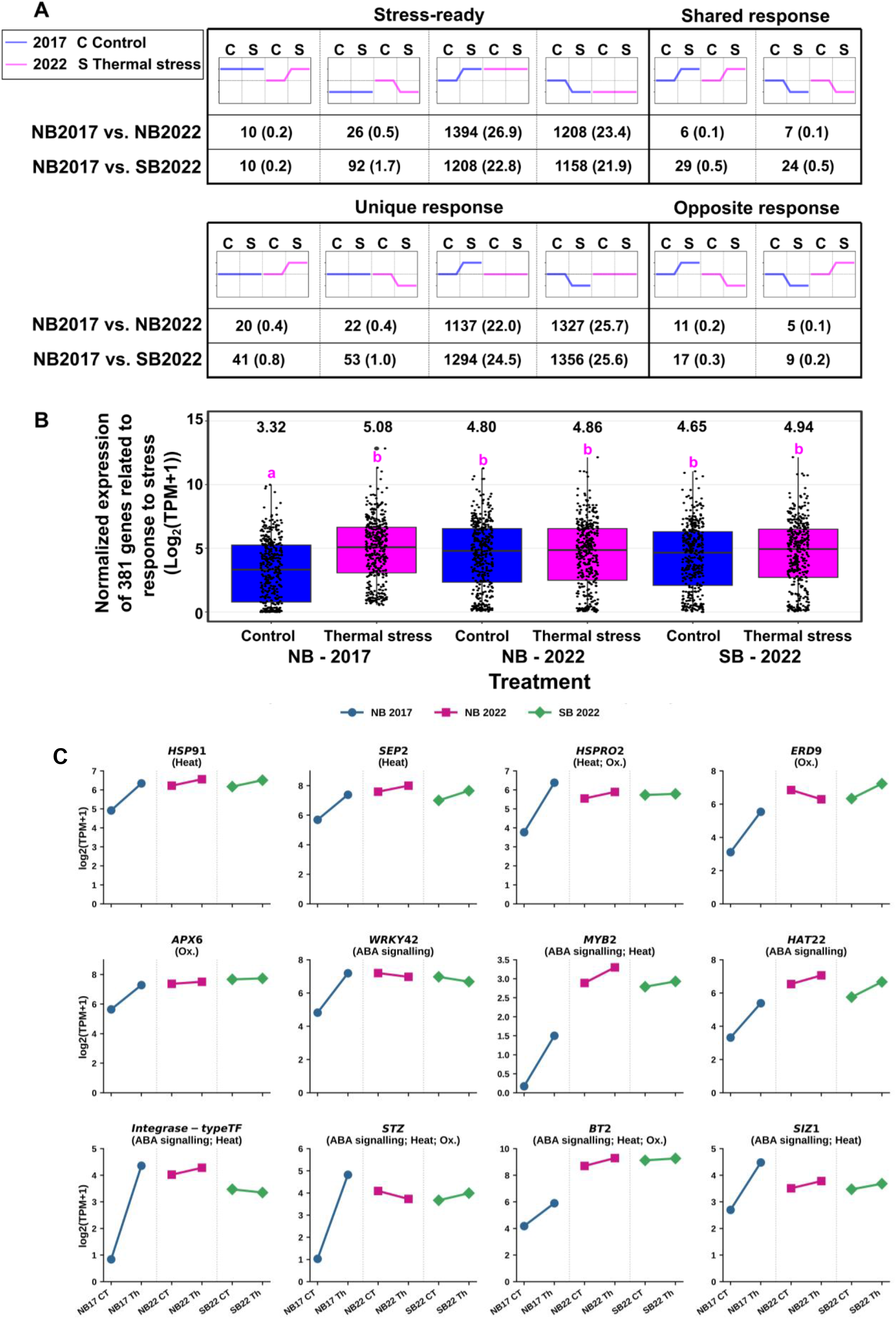
Transcriptomes from *Halophila stipulacea* plants collected in 2022 exist in a heat “stress-ready” state. (**A**) Gene expression modes in response to thermal stress in 2017 and 2022 plants. Blue (2017) and magenta (2022) lines indicate idealized expression patterns of homologous genes in each cohort under control and thermal stress, compared to control (dashed horizontal line). C, control; S, thermal stress; Numbers under graphs, no. of homologous gene pairs assigned to each response mode; Numbers in parentheses, percent of homologous gene pairs relative to the total number of homologs (NB2017 vs NB2022 [5173]; NB2017 vs. SB2022 [5291]) assigned to a response mode. (**B**) Normalized expression of 381 homologous DEGs related to “response to stress” (GO:0006950). whose expression was upregulated by thermal stress in 2017 plants vs. 2022 plants. For box and whisker/jitter plots, the median (thick black line and numbers above) and interquartile range are shown. Whiskers, the maximum/minimum range. Letters, significant differences (*p* < 0.05; Tukey HSD *post hoc* test, *n* = 3) between treatments based on a one-way ANOVA test of the median of normalized expression of genes. Prior to analysis, the homogeneity of variance assumption was checked using Levene’s test. Normality was assessed using the Shapiro-Wilk test. (**C**) Mean normalized expression profiles [log_2_(TPM + 1)] of representative stress-responsive genes selected from the 381 stress-related DEGs in (**B**), illustrating the “stress-ready” pattern across heat stress, oxidative stress, and ABA signalling categories. Expression values for each treatment represent averages across biological replicates (*n* =3) derived from Supplementary Table S7. Expression trajectories are shown under control (CT) and thermal stress (Th) for NB 2017 (blue circles), NB 2022 (magenta squares), and SB 2022 (green diamonds). Dotted vertical lines visually separate temporal (2017 vs. 2022) and spatial (NB vs. SB) cohorts. Gene names and functional stress categories (in parentheses) are derived from Supplementary Table S8.

Consistent with this evidence of a “Stress-ready” transcriptome in the 2022 plants, analysis of the expression of 381 DEGs associated with the GO-term “response to stress” (GO:0006950), which were upregulated by mesocosm-applied thermal stress in the 2017 plants revealed that these genes exhibited a “Stress-ready” mode of expression in both 2022 populations (Fig. 4B; Supplementary Table S7). Many of these genes encode proteins involved in plant abiotic stress responses. Notably, 49 genes were associated with heat stress, including late embryogenesis abundant proteins, heat shock proteins, heat shock factors, and molecular chaperones (Supplementary Table S8). An additional set of genes functioned in oxidative stress mitigation (peroxidases, glutathione- and ascorbate-related enzymes), while several were linked to abscisic acid signalling/response. This pre-emptive activation of protective networks is illustrated by representative genes spanning heat stress, oxidative stress, and ABA signalling categories (Fig. 4C; Supplementary Table S8). Heat-responsive genes such as *HSP91, SEP2* and *HSPRO2* showed strong induction under thermal stress in 2017 plants, yet in both 2022 populations their expression under control conditions already approached or exceeded the 2017 thermal-stress level, with little further change in response to thermal stress. Similar stress-ready trajectories were observed for oxidative-stress genes (*APX6, ERD9*) and multiple components of ABA signalling (*WRKY42, MYB2, HAT22, STZ, BT2, SIZ1* and an *Integrase-type transcription factor*). Collectively, these profiles demonstrate a coordinated shift toward constitutively primed defensive networks in the 2022 plants.

Taken together, our data support the notion that the transition from episodic to chronic thermal stress during the five-year period between 2017 to 2022 acted as a driver of molecular reprogramming resulting in a “stress-ready” thermal stress transcriptome that was associated with increased tolerance to ocean warming in 2022 *H. stipulacea* plants.

## DISCUSSION

This study provides the first longitudinal physiological and transcriptomic datasets documenting a five-year temporal divergence in thermal tolerance within a natural seagrass population, with transcriptomic shifts linked to corresponding changes in growth and photochemistry and associated with recorded local ocean warming and the transition from episodic to chronic thermal stress. These data suggest that at least one tropical species, *Halophila stipulacea*, is capable of rapid adjustment to warming. Plants collected in 2022 from the GoA maintained growth and photochemistry under experimentally applied thermal stress whereas conspecifics from the same site in 2017 suffered clear declines. This shift occurred within five years and was underpinned by an attenuated transcriptional response to thermal stress and a constitutively “stress-ready” transcriptome in which the expression of thousands of thermally responsive genes in the 2017 cohort was already up- or downregulated under control conditions in the 2022 plants. Furthermore, the increased thermal tolerance and the presence of a “stress-ready” transcriptome was not only restricted to conspecifics from NB but was also displayed in a separate population from SB.

The difference in thermal stress duration (21 days in 2017 versus 35 days in 2022) introduces the possibility that the attenuated 2022 response reflects a late-phase resolution of an acute stress response rather than a constitutive, “stress-ready” baseline. However, our data exclude a transient resolution model because the baseline control expression of these up-/downregulated “stress-ready” genes in 2022 plants closely matches their expression under thermal stress, and both metrics approximate the maximum acute thermal stress-mediated induction captured in the 2017 plants (Figs. 4, A and B; Supplementary Table S6). Similarly, variations in ambient seawater temperature at the time of collection could alter the effective experimental thermal delta. If 2022 field temperatures were elevated relative to 2017, a narrower differential (ambient to 32°C) could account for the diminished transcriptomic induction.However, *in situ* monitored ambient seawater temperatures at the collection sites were between 25-26°C across both years matching the control temperature used in the experiments, and thereby ruling out baseline field acclimatization effects.

Several non-mutually exclusive mechanisms could underlie the observed acquisition of a “stress-ready” transcriptome. Firstly, phenotypic plasticity could allow a single genotype to produce different phenotypes, including altered gene-expression profiles, in response to environmental cues (*36, 37*). Secondly, selection on standing genetic variation could favor alleles that promote constitutive regulation of stress-response genes; experimental warming of *Zostera marina* has already revealed substantial genotype-specific differences in thermal tolerance within a single population (*38*). Thirdly, stress-induced epigenetic modifications such as histone modifications or DNA methylation changes, could stably maintain the “stress-ready” state across generations. This notion is supported by thermal priming experiments on seagrasses that enhance thermal tolerance accompanied by changes in DNA methylation patterns and transcriptome reprogramming of stress-response genes (*39-41*). Indeed, it was recently demonstrated that in the pioneer seagrass, *Cymodocea nodosa*, thermal priming induces targeted gene-body hypomethylation in key stress-response genes - a modification that persists trans-clonally across ramets and could maintain the chromatin in a permissive state for rapid transcriptional activation upon re-exposure (*42*). In line with such mechanisms, the observation of the “stress-ready” transcriptome in 2022, a year with relatively few extreme-temperature days, (Fig. 2) is consistent with either selection on favorable alleles or/and the retention of permissive and repressive epigenetic marks. Whether the shift toward a “stress-ready” transcriptome represents genetic adaptation or neutral processes remains open. Space-for-time studies that have been performed require stringent correction for phylogeographic differentiation, with only a small fraction of transcriptomic divergence exceeding neutral expectations (*23*). In contrast, our longitudinal dataset ties transcriptomic reprogramming to the documented transition from episodic to chronic thermal stress. While this association is correlational, the rapid multi-year stabilization aligns with documented timescales for climate-driven clonal epigenetic inheritance and microevolution in plants (*41, 43, 44*), thereby strengthening the case for climate-driven adjustment.

Similar “stress-ready” transcriptomes have been observed in terrestrial halophytes adapted to persistent ionic stress (*34, 35*). In terms of terrestrial plant adaptation to thermal stress, the desert Brassicaceae *Anastatica hierochuntica*, although extremely heat-tolerant, does not possess a “stress-ready” transcriptome. Instead, it displays a hyperreactive transcriptome; one-third of its genes show stronger fold-change induction under heat stress than orthologs in the heat-sensitive model *Arabidopsis thaliana* (*30*). We suggest that the difference in thermal adaptation between terrestrial *A. hierochuntica* and aquatic *H. stipulacea* arises from habitat stability. Desert temperatures fluctuate dramatically (up to 18 °C diurnally in the Negev desert; Israeli Meteorological Services), thereby favoring a hyperreactive, on-demand transcriptional response. In the ocean, however, heat absorbed during warming events persists for days to weeks because of water’s high specific heat capacity, thereby creating a more chronic selective pressure akin to persistent salinity in halophyte habitats. This sustained pressure appears to favor a constitutively “stress-ready” transcriptome. This contention is supported by the finding that local selective forces can drive molecular reprogramming in *H. stipulacea*. Hence, GoA populations from anthropogenically-impacted (high nutrient load) sites possess lower stress-induced plasticity and constitutively expressed “stress-ready” genes (*22*). Accordingly, plants from the impacted site exhibit greater tolerance and resilience to combined thermal and nutrient stress than plants from the pristine site (*22, 24*). Our current study extends this pattern to a purely climatic driver across a documented warming trajectory, reinforcing the idea that contemporary ocean warming can impose sufficiently strong selection to produce detectable molecular shifts within a decade.

Our findings have direct implications for forecasting the responses of *H. stipulacea* to continued warming and may also shed light on its remarkable invasive success. This species has spread extensively throughout the Mediterranean and more recently expanded to where summer water temperatures routinely exceed those in many native-range coastal habitats (28, 45). The capacity for rapid transcriptomic reprogramming demonstrated here may partly explain this thermal versatility. If populations can adjust their stress responses within years rather than generations, range expansion into progressively warmer waters becomes more feasible.

However, such capacity may be species- or population-dependent. Recent studies show that different seagrass species vary markedly in their responses to prolonged warming, with some exhibiting acute sensitivity while others maintain performance (*46*).

Whether this rapid shift toward stress-tolerant phenotypes in response to ocean warming carries ecological trade-offs remains unanswered. Constitutive expression of stress-response genes may incur energetic costs that reduce growth rates, alter carbon balance, or modify interactions with herbivores, epiphytes, or associated fauna. Such trade-offs could reshape meadow structure and diminish the capacity of *H. stipulacea* beds to support reef-associated biodiversity under continued warming (*47*). Long-term field monitoring and multi-generational common-garden experiments will be essential to evaluate these potential costs and to determine whether similar adaptive capacity exists in other seagrass species. If rapid transcriptomic preparedness proves widespread, it could substantially alter current forecasts of seagrass persistence in warmer seas.

## MATERIALS AND METHODS

### Study area and plant collection

The northern Gulf of Aqaba (GoA, known in Israel as the Gulf of Eilat) is an oligotrophic ecosystem with clear water [light attenuation coefficient Kd _[400 to 700 nm]_ = 0.054 (*48*)] and a long water residence [3-8 years; (*49*)] due to its unique geographical location (i.e. it is surrounded by arid regions and it has limited water exchange with the Red Sea due to the Straits of Tiran). Here, the tropical seagrass *Halophila stipulacea* is the only seagrass species and it forms monospecific meadows on soft sediments ranging between 2-50 m depth between local coral reefs (*26*).

For the current study, *H. stipulacea* populations from two different sites in the northern GoA were selected (Fig. 1, A and B). The North Beach (NB; 29°32’46.7”N 34°57’50.5”E) site is surrounded by the center of the city of Eilat with many hotels and busy urban areas. In contrast, the South Beach (SB; 29°29’34.2”N 34°54’21.6”E) site is located at the southern edge of the city, far away from most hotels and urban activities. Long-term monitoring data have shown that throughout the year: (i) the water is less clear at the NB site than at the SB site; (ii) the level of nutrients in the water column (e.g. nitrate, nitrite, ammonium and phosphate) at the NB site is much higher than nutrients at the SB site (*27*). Therefore, the NB site is considered an impacted site while the SB site represents a pristine site. At the NB site, *H. stipulacea* meadow depths range from 8 m to 12 m along a low bathymetric slope (∼5°), with low coral coverage (*26*). On the other hand, at the SB site, *H. stipulacea* meadows can be found at a depth range of 6 m to over 50 m along a steep bathymetric slope (∼18°), with high coral coverage (*26*).

On October 3^rd^ 2017, plants were collected by free diving at a depth of 4-5 m from the NB site (*28*). For the current study, plants were collected on the 28^th^ of November 2022 by SCUBA diving at a depth of 10-12 m from both the NB and SB sites. To avoid collection of the same clone twice, each *H. stipulacea* plant was collected 5-10 m away from each other.

Collected plants were kept in zip-lock plastic bags filled with natural seawater and quickly transported in cooler boxes to the seagrass mesocosm facility at the Dead Sea and Arava Science Centre (Hatseva, Israel) within less than 2 h post-collection. At the time of sample collection, seawater temperatures were between 25-26 ° C at both sites for both years (*28*; this study).

### Mesocosm experimental setup

Six aquaria (40 x 40 cm, 33 cm height) were filled with natural sediment, collected from the shores of Eilat, to a height of ∼7 cm and 40 L of artificial seawater (Red Sea Salt, Israel). To standardize the experiment, plants with a similar number of shoots (4-5 shoots) were selected and 9-10 plants from the NB and SB populations (i.e. 18-20 plants in total) were planted in each aquarium (Fig. 1C). Plants from the two populations were separated by a plastic divider, which allowed them to grow under the same conditions while avoiding any undesired population mixing. The position within each aquarium (i.e. left or right) of the NB and SB populations were assigned randomly between aquaria.

Before the start of the experiment, plants were allowed to acclimatize to the mesocosm conditions for 4 weeks. Water temperature was kept at 26 °C which matches the maximum annual seawater temperature occurring summer in the Gulf of Eilat (*27*). A 13 h/11 h light/dark photoperiod (similar to summertime) was applied starting from 08:00, where irradiance was progressively increased to reach a maximum level of ∼200 μmol photons m^-2^ s^-1^ at the canopy height (i.e. similar to the *in situ* irradiance at 10 m depth at the NB site during the summer) at 12:00 noon. Maximum irradiance was maintained for 3 h before gradual reduction to darkness at 21:00. The salinity level was maintained at 40 PSU by adding distilled water to compensate for evaporation, and approximately 1/4 of the water from each aquarium was renewed weekly to sustain the water quality.

After acclimation, the 6 aquaria were assigned to 2 different treatments (*n* = 3 aquaria for each treatment) as follows: Control, (26 °C); Thermal stress (32 °C) (Fig. 1D).

For the control aquaria, water temperature was kept at 26 °C, similar to the acclimation period (Fig. 1D, right panel). For the thermal stress aquaria, water temperature was ramped up from 26 °C to 32 °C over the course of a week (1 °C per d), and kept at 32 °C for 4 weeks (Fig. 1D, right panel).

For the duration of the experiment, salinity and light conditions of all aquaria throughout the experiment were maintained at the same level as during the acclimation period. Water temperature was measured every hour and controlled automatically using a ProfiLux 3 Controller (GHL, Germany).

### Growth and photo-physiological measurements

Plant growth, measured as the total number of shoots per plant, was measured weekly throughout the experiment. To standardize the experiment before application of stress (i.e. 0 d of stress), plants were selected that possessed exactly six shoots per plant, including one apical shoot. For each aquarium and each population, 3 plants were randomly selected and marked for the measurement of the total number of shoots. Subsequently, these three measurements were averaged into a single biological replicate (*n* = 3 per treatment per population).

Photo-physiological responses of the plants, including maximum potential quantum efficiency (Fv/Fm) and effective quantum yield (ΔF/Fm’) of Photosystem II, were measured weekly using a diving-PAM fluorometer (Walz, Germany) following the methodology described in (*50*). Fv/Fm was measured on whole-night dark-adapted plants (around 06:30-07:30 before lights-on) while ΔF/Fm’ was determined on light-adapted plants at the highest irradiance (around 13:00-14:00). For each aquarium and each population, two plants were randomly selected for the measurements. Data from these two plants were averaged and considered as one replicate (*n* = 3 per treatment per population).

### Estimation of average rise in Maximum Daily Seawater Temperature (MDST) over time

Maximum Daily Seawater Temperature (MDST) data in the Gulf of Eilat for the period 2006-2022 were obtained from the Israel National Monitoring Program in the Gulf of Eilat (www.meteo-tech.co.il) (Fig. 2). To quantify the magnitude of ocean warming, we calculated the average MDST rise relative to a baseline year using the following formula:

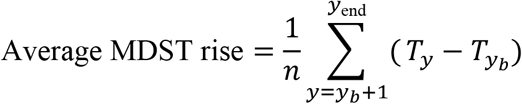

where *T*_*y*_ is the MDST in year *y, y*_*b*_ is the baseline year, and *n* is the number of years after the baseline.

Using 2006 as the baseline year, the average MDST rise from 2007 to 2022 was 0.57 °C. Using 2017 as the baseline year, the average MDST rise from 2018 to 2022 was 0.66 °C.

### Transcriptome sequencing and analysis

#### RNA sample collation, extraction and sequencing

In 2017, we conducted a mesocosm experiment to study the responses of native and invasive populations of *H. stipulacea* to predicted ocean warming (*28*). In comparison to the current study with plants from 2022, the 2017 experiment applied the same control and thermal stress levels (i.e. 26 °C and 32 °C, parallel samples for 21 d) and the samples were collected from the same *H. stipulacea* meadow at the NB site. The duration of stress was longer in the current study (i.e. 35 d of stress). RNA samples isolated from samples from the 2017 experiment were preserved in a - 80 °C freezer.

For both 2017 and 2022 *H. stipulacea* plants, leaves from the 2^nd^ youngest shoot were collected from a randomly selected plant from each population and each aquarium of the control and thermal stress aquaria (*n* = 3 per treatment per population). Immediately after collection, the leaves were gently cleaned from epiphytes and preserved in RNAlater^®^ (Thermo Fisher Scientific, USA). Samples were first stored at 4 °C overnight before being stored at -20 °C until RNA extraction.

For the 2017 plants, total RNA was extracted by TRIzol™ Reagent (Invitrogen; Thermo Fisher Scientific, Waltham, MA, USA) following the manufacturer’s instructions (*28*). Extracted total RNA was then treated with RNase-free DNase I (EN0525, Thermo Fisher Scientific). For the 2022 plants, total RNA was isolated using a Plant/Fungi Total RNA Purification Kit including on-column DNase I treatment (Norgen Biotek Corp., Canada) according to the manufacturer’s instructions. To prevent oxidation via phenolic compounds, 2% (w/v) polyvinylpyrrolidone-40 (PVP) was added to the lysis solution. RNA purity was assessed with a NanoDrop® ND-1000 spectrophotometer (Thermo Fisher Scientific, Waltham, MA, USA) and its integrity was checked on a 1% agarose gel electrophoresis.

RNA samples were sent to the Roy J. Carver Biotechnology Centre, University of Illinois (Urbana-Champaign, IL), where RNA integrity was assessed on an Agilent Bioanalyzer prior to library preparation and sequencing on an Illumina NovaSeq 6000 system to generate 150 nt paired-end reads.

#### Bioinformatics, differential expression analysis and functional annotation

Raw sequencing reads (Supplementary Table S2) were first quality-verified using FastQC v0.12.1 (*51*). Sequencing reads were filtered from rRNA using SortMeRNA v4.3.6 (*52*) and then mapped to the *H. stipulacea* genome (BioProject PRJNA1140917) using STAR v2.7.11a (53).

The average mapping rate across all samples was 87%. Differential expression was evaluated using DESeq2 v1.34.0 with adjusted *p* value < 0.05 (*54*; Supplementary Table S4). The PCA plot was visualized on rlog-transformed counts. For assignment of genes to modes of expression (“Stress-ready”, “Shared response”, “Unique response”, “Opposite response”), log_2_ fold-change in expression was compared between homologous genes of 2017 and 2022 plants that had been called as DEGs by DESeq2 (*22*). Differences in basal (control) levels of expression were assessed by comparing transcripts per kilobase million (TPM). For log_2_ fold-change in expression and TPM values for all DEGs, see Supplementary Table S6.

GO-term enrichment of DEGs was performed using BiNGO v3.0.3, available in Cytoscape v3.10.0 (*55, 56*), on *H. stipulacea*’s whole genome functional annotation using a hypergeometric test with FDR correction and adjusted *p* value < 0.05 for over-represented GO terms (*57*). Clustering of enriched GO-terms was performed using GOMCL v0.0.1 (*58*). Visualization of the clustered groups was performed in Cytoscape v3.10.0 (*55*).

## Acknowledgments

The authors would like to thank Beery Yaakov for technical assistance, and the team at the Roy J. Carver Biotechnology Center, University of Illinois Urbana-Champaign for their usual excellent sequencing services.

## Author contributions

Conceptualization: G.W., and S.B.Investigation: H.M.N., N.L.L, and M.K. Data analysis: H.M.N., G.W., and S.B. Writing: S.B., H.M.N., and G.W. Supervision: G.W., and S.B.

## Data and materials availability

The *Halophila stipulacea* genome data are available under BioProject accession PRJNA1140917 at the NCBI BioProject database. The transcriptome data generated in this study have been deposited in the NCBI BioProject database under BioProject accession PRJNA1180108. All other data are available in the manuscript or the supplementary materials.

## Supplementary Materials

Materials and Methods

Supplementary Figure S1

References (*47-57*)

Supplementary Tables S1 to S8

