## Supplementary Figure S1 for "Decadal warming is linked to thermal tolerance and transcriptome reprogramming in a seagrass"

**The PDF file includes:**

Supplementary Figure S1

**Other Supplementary Materials for this manuscript include the following:**

Supplementary Tables S1 to S8

**A**

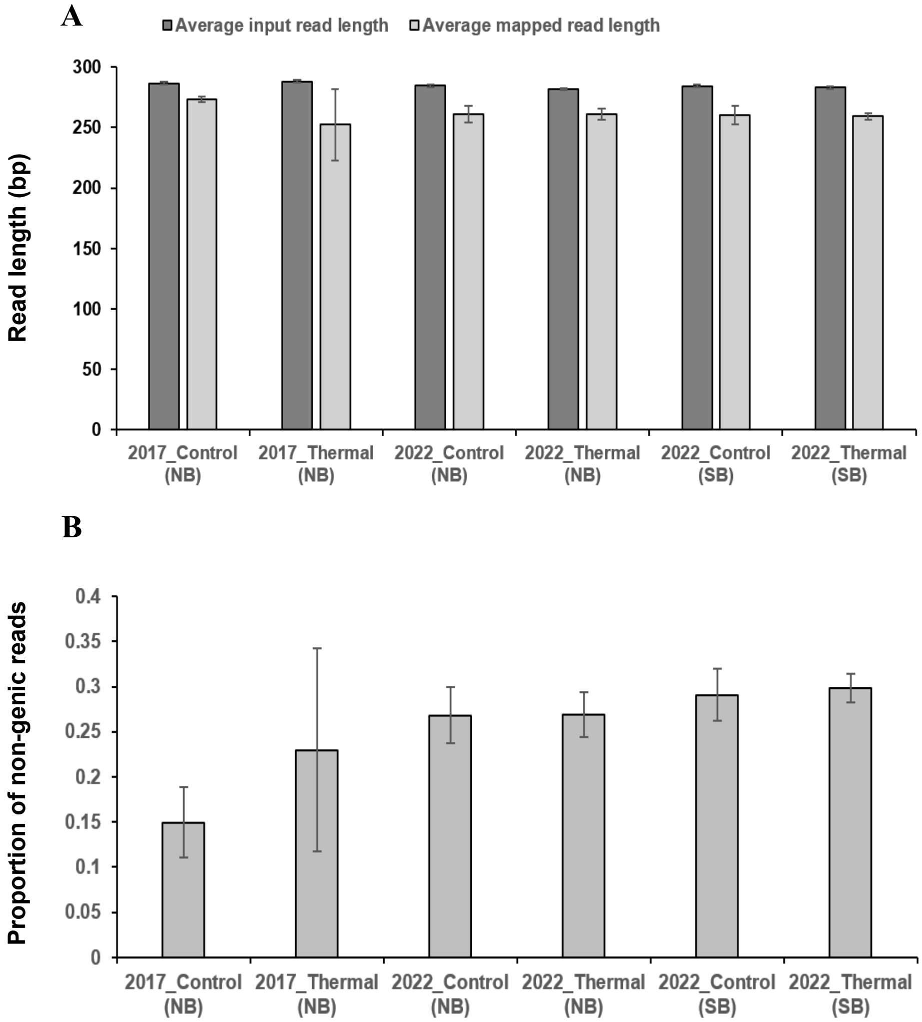


**Supplementary Figure S1. RNA sequencing quality metrics demonstrate comparable RNA quality between the 2017 and 2022 samples. (A)** Average input read length and average mapped read length for RNA-seq libraries generated from *Halophila stipulacea* plants collected in 2017 and 2022 under control (CT) and thermal stress (Thermal) conditions (n = 3). See materials and methods for experimental details. **(B)** Fraction of sequencing reads mapped to non-genic regions. Quality control metrics were obtained during RNA-seq alignment using STAR v2.7.11a (*52*).

RNA-seq quality metrics indicate that RNA quality was comparable between samples collected in 2017 and 2022 despite the approximately five-year longer storage of the 2017 RNA prior to library preparation. Average input and mapped read lengths were highly similar across all sample groups, indicating no evidence of increased RNA fragmentation in the older samples. Likewise, the proportion of reads that mapped to non-genic regions was comparable between years, demonstrating similar mapping quality and transcriptome representation. These observations are consistent with the RNA integrity measurements (RQN values) and sequencing yields (Supplementary Table S3), which likewise showed no reduction in RNA quality in the 2017 samples. Together, these independent quality-control metrics demonstrate that long-term storage of the 2017 RNA did not measurably compromise RNA integrity or RNA-seq library quality, indicating that the transcriptomic differences observed between the 2017 and 2022 populations are unlikely to be attributable to differential RNA preservation.

**Supplementary Table S1.**
Two-way ANOVA for growth and photo-physiology.

**Supplementary Table S2.**
Raw read counts with GO term annotations for all expressed genes.

**Supplementary Table S3.**
RNA integrity metrics (RQN) and sequencing depth.

**Supplementary Table S4.**
Differentially expressed genes (DEGs) in response to thermal stress with gene ontology-terms.

**Supplementary Table S5.**
Gene ontology-term enrichment of up-/downregulated DEGs in response to thermal stress.

**Supplementary Table S6.**
Assignment of genes to idealized expression response-modes.

**Supplementary Table S7.**
Expression of genes related to gene ontology-term "response to stress" (GO 0006950).

**Supplementary Table S8.** Heat, oxidative stress, and ABA signaling gene categories derived from Supplementary Table S7.
